# Neural overlap in item representations across episodes impairs context memory

**DOI:** 10.1101/125971

**Authors:** Ghootae Kim, Kenneth A. Norman, Nicholas B. Turk-Browne

## Abstract

We frequently encounter the same item in different contexts, and when that happens, memories of earlier encounters can get reactivated in the brain. Here we examined how these existing memories are changed as a result of such reactivation. We hypothesized that when an item’s initial and subsequent neural representations overlap, this allows the initial item to become associated with novel contextual information, interfering with later retrieval of the initial context. That is, we predicted a negative relationship between representational similarity across repeated experiences of an item and subsequent source memory for the initial context. We tested this hypothesis in an fMRI study, in which objects were presented multiple times during different tasks. We measured the similarity of the neural patterns in lateral occipital cortex that were elicited by the first and second presentations of objects, and related this neural overlap score to source memory in a subsequent test. Consistent with our hypothesis, greater item-specific pattern similarity was linked to worse source memory for the initial task. Our findings suggest that the influence of novel experiences on an existing context memory depends on how reliably a shared component (i.e., same item) is represented across these episodes.

## Introduction

Our experience is highly repetitive, with the same objects appearing repeatedly over time and often in different contexts. For example, we might move a piece of furniture to many different apartments, see somebody from work at the grocery store, or look for our car in various parking lots. How does experiencing a familiar item in a novel context affect preexisting memories of the item and its prior contexts?

It has long been known that memory for the initial context in which an item was experienced can be impaired by a later encounter of the item in a new context. Such *retroactive interference* has been widely investigated using the AB/AC paradigm (McGovern, 1964; Postman & Underwood, 1973; Richter, Chanales, & Kuhl, 2016). In this paradigm, participants learn an episode with components A and B, then another episode with components A and C. As in a previous example, we might encounter a colleague at the grocery store (item A in context C), with whom we previously chatted in the office (item A in context B). Because of the shared component A, learning AC can trigger retrieval of the previously learned AB memory. How does this memory reinstatement relate to retroactive interference (i.e., forgetting of B and/or AB)? One prominent account is that reactivation of a prior context B during later AC learning builds resistance to interference, leading to better subsequent retrieval of the initial context B when cued with A (Koen & Rugg, 2016; Kuhl, Shah, DuBrow, & Wagner, 2010).

Here we investigate a different, though not mutually exclusive, account of how memory reinstatement relates to retroactive interference. We focus on the fact that mental representations of an item can differ over time even when we putatively experience the “same” item. In the AB/AC paradigm, for example, this would correspond to variance in the extent to which the representation of A during AC learning is the same as the representation of A during the prior AB learning. Although neural overlap across repeated presentations of an item can be associated with better memory for that item (Ward, Chun, & Kuhl, 2013; Xue et al., 2010), it is unknown how this overlap affects memory for previously formed item-context associations.

We hypothesize that retroactive interference occurs when the same *item* representation is reinstated across episodes with different contexts. Specifically, reinstatement of the item representation engaged by the initial processing of A (from AB learning) during AC learning allows this prior item to become associated with the novel context (C), which interferes with later retrieval of the initial context B. In contrast, if the representation of A during AC learning differs from that of the prior AB episode, memory of the initial context B might be less affected by retroactive interference. In short, we predict a negative relationship between item-specific representational overlap and subsequent source memory for the initial context. Note that this is the opposite of the positive relationship found for context (rather than item) reinstatement by Koen and Rugg (2016) and Kuhl et al. (2010), where more context information leads to greater initial source memory.

To test this hypothesis, we presented objects (A) sequentially during two different orienting tasks (B and C). These tasks served as the contexts to which the items could be bound (Johnson, Kounios, & Nolde, 1997). Using fMRI, we measured pattern similarity for a given item across the two task contexts in the lateral occipital cortex (LOC), which is thought to represent the visual features of objects (Grill-Spector, Kourtzi, & Kanwisher, 2001). We then related these item-wise pattern similarity scores to subsequent source memory for the initial task. In addition to testing for the hypothesized negative effect of item reactivation, we also tested for the positive effect of context reactivation observed in previous studies (Koen & Rugg, 2016; Kuhl et al., 2010). Such a dissociation would provide strong evidence that item and context reactivation have differential effects on source memory. Consistent with our main hypothesis, we found a negative relationship between item reactivation and subsequent source memory: greater item-wise pattern similarity was associated with worse source memory for the initial task.

## Materials and methods

### Overview

This study consisted of four phases. In an initial encoding phase (phase 1), participants were exposed to a sequence of object images and performed one of two orienting tasks (artist or function task). These tasks served as the initial context to which the object items could be bound during the encoding phase. In the item repetition phase (phase 2), half of these objects were presented again while participants performed a new orienting task (organic task). We measured how reliably the initial representations of the items were reinstated in this phase by calculating item-specific pattern similarity between the initial and repeated presentations. In the memory test phase (phase 3), source memory for the initial task (i.e., artist or function) was measured and related to the item-specific pattern similarity scores calculated in the second phase. A final task localizer phase (phase 4) was used to generate template neural patterns for each task, which were used to measure task representations during phases 1 and 2 for secondary analyses.

### Participants

Thirty-two adults (14 women, all right-handed, mean age 21.88 years) participated for monetary compensation. All participants had normal or corrected-to-normal vision and provided informed consent. The Princeton University IRB approved the study protocol.

### Stimuli

Participants were shown color photographs of natural and manmade real-world objects. Stimuli were displayed on a projection screen behind the scanner bore, viewed with a mirror on the head coil (subtending 8.8 × 8.8°). Participants fixated a central dot that remained onscreen throughout.

### Procedure

Participants completed one scanning session with four phases: initial encoding, item repetition, source memory test, and task localizer. During the initial encoding phase (Fig. 1A), participants viewed a series of objects that were randomly assigned to one of two orienting tasks: How easy would it be to draw the object? (artist task) or How useful is the object? (function task). Participants responded on a 4-point scale (artist/function): 1 = very easy/very useless, 2 = easy/useless, 3 = hard/useful, 4 = very hard/very useful. We used these tasks because previous studies have shown that they are highly decodable with fMRI (Johnson, McDuff, Rugg, & Norman, 2009; Koen & Rugg, 2016; McDuff, Frankel, & Norman, 2009). Four runs of encoding were collected, and each run contained 2 blocks of objects from each of the two tasks (the order of the 4 blocks was randomized). The task was instructed with a cue at the beginning of the block (e.g., “Artist task”). Each object stimulus was presented for 1 s, followed by a blank interval of 2 s. There were 12 trials per block (36 s duration), followed by 15 s of rest. The total duration of each run was 3 min 42 s.

**Figure 1.**
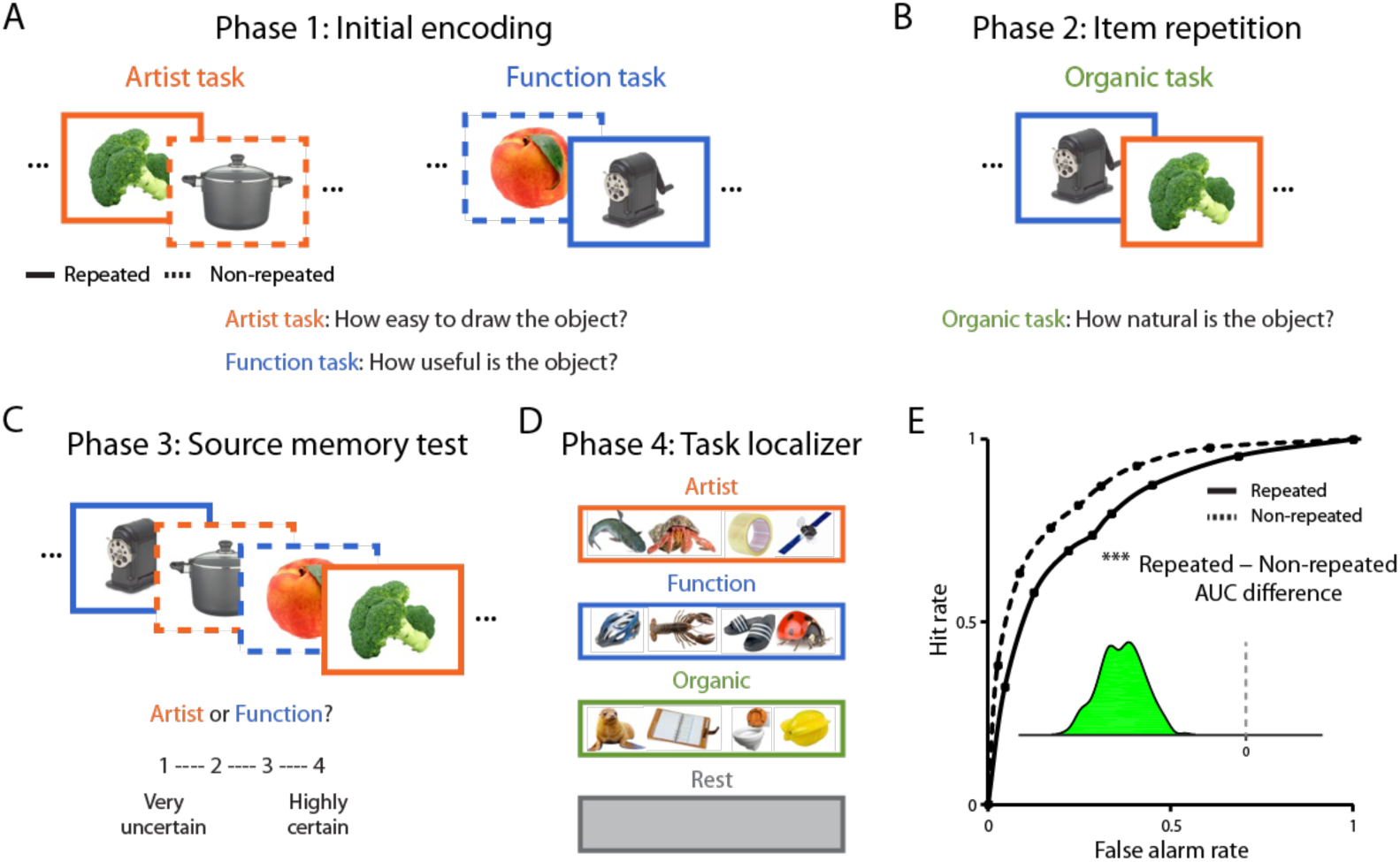
Experimental design and behavioral results. (A) During initial encoding, object images were randomly assigned to one of the two orienting tasks (artist and function tasks). (B) In the item repetition phase, half of objects from the first phase were presented again while participants performed a third (organic) task. (C) In the subsequent source memory test, judgments were collected about which task had been performed first on each object, both for objects presented twice (repeated condition) and objects presented once (non-repeated condition). (D) In the task localizer, a new set of objects was presented in each of the three tasks to define task-specific neural activity patterns. (E) The area under the curve (AUC) of source memory judgments was calculated; lower AUC indicates worse memory, and so memory for the first task was worse in the repeated vs. non-repeated condition. The inset plot depicts the sampling distribution of the repeated minus non-repeated AUC difference from random-effects bootstrap resampling of participants. Almost all resampled AUC differences were below zero (green area), indicating a reliable retroactive interference effect. *** *p* < .001.

In the item repetition phase (Fig. 1B), half of the objects from each task in the initial encoding phase (i.e., 48 objects for each of artist and function) were presented again, and participants determined how organic the object was on a 4-point scale: 1 = very artificial, 2 = artificial, 3 = natural, 4 = very natural. Each of the 96 objects was presented for 1 s, followed by a blank interval of 3.5 s. All stimuli were presented in a single run without a rest period, lasting 7 min 33 s.

The source memory test (Fig. 1C) came as a surprise to participants. It contained the 96 objects presented in both the encoding and repetition phases (repeated condition) and the 96 objects shown only in encoding phase (non-repeated condition). Participants were instructed to discriminate which task had been performed on each object during the initial encoding phase (i.e., artist or function), and then to report their confidence level on a 4-point scale: 1 = very unsure, 2 = unsure, 3 = sure, and 4 = very sure. Each object was presented for 6 s, though participants were encouraged to respond within 5 s. If they failed to respond on a given trial, the object was omitted from later analyses (around 3% of total trials).

After the memory test, participants completed three runs of a functional localizer (Fig. 1D), in which new object images were presented in one of the three tasks: artist, function, and organic. Each run contained 6 blocks, with 2 blocks from each of the three tasks in a random order. Each object was presented for 1 s, followed by a blank interval of 2 s. There were 12 trials per block (36 s duration). Each block was followed by 15 s of fixation, which was treated as a baseline “rest” category. Total run duration was 5 mins 24 s.

### Behavioral analysis

We measured memory performance by dividing responses from the source memory test into eight levels of confidence: 4 = very sure “artist” to -4: very sure “function”. These judgments were quantified using receiver operating characteristic (ROC) analyses (Macmillan & Creelman, 2005; Green & Swets, 1966). For each of the repeated and non-repeated conditions, we created an ROC curve across the eight confidence levels and calculated the area under the curve (AUC). Calculating these curves precisely requires a substantial amount of data, and thus we pooled trials across participants beforehand. We assessed the reliability of the AUC difference between conditions across participants using a bootstrapping approach in which entire participants were resampled with replacement 1,000 times (Efron, 1979), and AUC was computed (for each resampling) based on the trials pooled across all resampled participants. This provided a population-level confidence interval (CI) for each effect, and also allowed for null hypothesis testing based on the proportion of bootstrapped samples in which the effect was reversed.

### Data acquisition

Experiments were run with the Psychophysics Toolbox (http://psychtoolbox.org). Neuroimaging data were acquired using a 3T MRI scanner (Siemens Skyra) with a 16-channel head coil. A scout anatomical scan was used to align axial functional slices. Functional images covering the whole brain were acquired with a T2* gradient-echo EPI sequence (TR = 1.5 s; TE = 28 ms; flip = 64°; iPAT = 2; matrix = 64×64; slices = 26; thickness = 4 mm, resolution = 3×3 mm). High-resolution (MPRAGE) and co-planar (FLASH) T1 anatomical scans were acquired for registration, along with field maps to correct B0 inhomogeneities.

### Preprocessing

fMRI data were preprocessed with FSL (http://fsl.fmrib.ox.ac.uk). Functional scans were corrected for slice-acquisition time and head motion, high-pass filtered (128 s period cutoff), spatially normalized (5mm FWHM), and aligned to the middle volume.

### Selection of ROIs

We defined ROIs for object processing (LOC) and for task processing. LOC was defined anatomically from the Harvard-Oxford cortical atlas in FSL and transformed into each participant’s space. The task ROI was defined in two steps: (1) We picked voxels selective to each of the three tasks (artist, function, and organic) by performing a general linear model (GLM) analysis of the localizer, with regressors for each task and rest. We ran three contrasts (artist vs. others, function vs. others, and organic vs. others) and selected voxels whose absolute *z*-values were above 2.3 (*p* < .01). (2) We then took the union of the surviving voxels of each contrast and defined a mask on an individual-subject basis.

### Measuring item reactivation

We measured how reliably the initial representation of each item was reinstated in the second phase by calculating the Pearson correlation of the patterns of activity elicited in the LOC on the initial and repeated presentations, 4.5 s after stimulus onset.

### Measuring task information

We measured the amount of task information in the first phase (to index task encoding) and in the second phase (to index task reactivation). Based on the localizer, we defined an activation template for each of the initial tasks (i.e., artist and function) over the voxels of the task mask by averaging patterns from the corresponding task (e.g., patterns from the artist task). Patterns of activity from the first and second phases were then correlated with the templates. We subtracted the correlation with the irrelevant task template from that of relevant template to get a difference score (e.g., when measuring task reactivation in the second phase, the artist template was relevant and the function template irrelevant for items whose initial task was artist).

### Relating neural measures to source memory

We measured the relationship between each of the neural measures (item reactivation, task encoding, and task reactivation) and subsequent source memory. For each neural measure, we divided trials into high vs. low pattern similarity groups based on a median split, and computed an ROC curve within each split. We then measured the area under each of the ROC curves (AUC), which reflects sensitivity of source memory (i.e., larger AUC is associated with better source memory). To calculate the effect of pattern similarity, we computed the difference between the high-pattern-similarity AUC and the low-pattern-similarity AUC. The sign of this AUC difference represents the direction of the relationship between the neural measure and memory: positive sign = positive relationship (i.e., better memory when pattern similarity is higher) and negative sign = negative relationship (i.e., worse memory when pattern similarity is higher). We performed this analysis by pooling trials across participants to measure a reliable relationship, and we assessed the population-level reliability of the result using a bootstrap procedure where we resampled participants with replacement. When running this analysis, we standardized pattern-similarity values within each participant before pooling them together; this ensures that any relationship we observe between a pattern similarity and subsequent memory reflects within-participant variance as opposed to across-participant variance.

### Simulations of approaches for measuring item-specific reactivation

Our main hypothesis concerns the relationship between the reactivation of an item’s representation and subsequent source memory. To test this, we needed a way of measuring reactivation of item features. Our basic approach (as described above) was to compute pattern similarity in LOC for the two presentations of each stimulus in the initial encoding (phase 1) and item repetition (phase 2) phases, respectively, and then to correlate this measure with source memory for the task from initial encoding (e.g., phase 1). However, this analysis is complicated by the fact that LOC pattern similarity is potentially influenced by two factors — reactivation of item features and reactivation of generic artist or function task features — which could influence memory in different (potentially opposing) ways. That is, reactivation of the *item* representation could lead to retroactive interference (for reasons described in the Introduction) whereas reactivation of *task* features could boost subsequent source memory (as in Koen & Rugg, 2016; Kuhl et al., 2010). For this reason, it was essential to use an analysis procedure that could separate the effects of item vs. task reactivation.

To compare different procedures, some that we devised and others used in the literature (Kim et al., 2014; Koen & Rugg, 2016), we ran simulations of two situations: (1) where subsequent memory was affected by item reactivation but not task reactivation, and (2) where subsequent memory was affected by task reactivation but not item reactivation. Our goal was to find an analysis procedure that would report a positive result only in the first situation (i.e., item reactivation is driving subsequent memory). The simulation results are summarized here, and the simulation methods are described in fuller detail in the Supplementary Material.

#### Approach 1: same-item minus different-item pattern similarity

Similar to our study, Koen and Rugg (2016) investigated the effects of item-specific pattern reactivation on source memory. To measure item-specific reactivation (while excluding task reactivation), they computed pattern similarity for pairs of the same item (e.g., A-first-presentation to A-second-presentation) and subtracted out the average pattern similarity of that item with other items encoded with the task (e.g., A-first-presentation to B-second-presentation, B-first-presentation to A-second-presentation, where A and B were encoded in the same task). They then related this difference measure to source memory.

Intuitively, one might think that comparing A to other items (B, C) encoded with the same task would control for task reactivation. However, in our simulations, we found that this method yielded significant results both when memory was driven by item reactivation *and* when memory was driven by task reactivation but not item reactivation (*ps* <.001). This result can be explained by the fact that same minus different pattern similarity controls for the *average* level of task reactivation, but it does not fully control for trial-by-trial variability in task reactivation (see Supplementary Material). Given that this analysis does not decisively discriminate item and task reactivation effects on memory in our simulations, we explored other approaches.

#### Approach 2: permutation analysis

We previously used permutation analysis to track reactivation of item features (Kim at al., 2014). This involves scrambling the pairings of items (across phase 1 and phase 2) 1,000 times and, for each scramble, re-calculating pattern similarity and its relationship to memory. A *z*-score of the original effect (based on the intact pairings) with respect to this null distribution can be calculated.

This analysis produced different results across the two simulations (situations A and B). When item reactivation drives memory, the original effect was reliably greater than the permuted effects (*p* < .001). In contrast, when task reactivation drives memory, the original effect was not different from the permuted effects (*p* = .53). This pattern of results confirms that the permutation analysis can identify relationships between item feature reactivation and memory and is not misled by trial-by-trial variance in task reactivation.

#### Approach 3: regression + permutation analysis

Although the permutation analysis handles the case where task reactivation in the second phase varies across items, it can be misled if task activation at encoding (for a given item) is correlated with task reactivation for that item at retrieval. That is, if a task is highly active during encoding, it might be reactivated more during the item repetition, and permuting the item pairings will eliminate this, giving the appearance of item-specific information. Indeed, when we simulated this situation (i.e., where reactivation of task features but not item features is correlated with subsequent memory, and task activation at encoding for a given item is correlated with task reactivation), the permutation test yielded a significant result (*p* < .001). The fact that the permutation test can show a significant result when (in the simulated data) there is no actual relationship between item feature reactivation and subsequent memory indicates that this test is also unsuitable for our purposes.

To address this issue, we adopted a different approach of regressing out the localizer template activity patterns for each task from every item’s representation prior to calculating pattern similarity and its relationship to source memory. After removing task information in this manner on a trial-by-trial basis, our simulation where item-specific reactivation drives memory survives the permutation test, but both forms of the simulation where task reactivation drives memory (i.e., where there is item-wise variance in task reactivation alone, and where there is also correlated item-wise variance in task encoding and task reactivation) both fail for the first time (*ps* > .37).

Having confirmed the validity of regression + permutation analysis, we applied it to our data in three steps: First, we regressed out the localizer template for each task from the corresponding patterns in the first and second phases (e.g., for an item whose initial task was artist, we regressed out the artist template from both patterns for that item). We then measured item-wise pattern similarity in the residual patterns, and related it to source memory by measuring the AUC difference for items with high vs. low pattern similarity. Second, we performed a permutation analysis by scrambling the original pairings of first and second phase trials within each task and participant (e.g., permuting items whose initial task was artist). For each of 1,000 permutations, we re-calculated pattern similarity and related it to memory. Finally, a *z*-score of the original, task-residualized relationship (AUC difference score) based on the intact pairings was calculated with respect to the null distribution of AUC differences.

## Results

### Subsequent source memory behavior

Overall, participants successfully discriminated the correct and incorrect initial task (mean *A*′ = 0.56, bootstrap *p* < .001). Consistent with the idea that item repetition increases retroactive interference, source memory for the initial task was lower for repeated compared to non-repeated items (AUC difference = -0.07, CI = [-0.09, -0.04], bootstrap *p* < .001; Fig. 1E).

### Initial pattern similarity results

We first confirmed that LOC activation patterns contained information about items (Cichy, Chen, & Haynes, 2011; Eger, Ashburner, Haynes, Dolan, & Rees, 2007), by showing that pattern similarity between the first and second phases was greater when the item was the same vs. when two different items from the same task were compared (bootstrap *p* < .001). To address the possibility that this pattern might reflect item-wise variance in task activation rather than item-specific information, we performed a two-step permutation analysis (see Methods). First, we regressed out generic task information from patterns in the first and second phases, and recalculated the same-item pattern similarity. Second, we permuted the original item pairings across the first and second phases and re-measured the item-wise pattern similarity. Finally, we tested whether the same-item pattern similarity from intact pairings is greater than the item-wise pattern similarity scores of permuted pairings. Indeed, the original same-item pattern similarity was reliably more positive than a null distribution of permuted pattern similarity scores (average *z*-score = 5.36, CI = [1.65, 9.29], bootstrap *p* < .001 Fig. 2A). We next confirmed that activation patterns from the task masks (which were defined from the localizer) contained task information in the first phase. Indeed, across trials, pattern similarity was higher for the relevant (i.e., currently-being-performed) task template than the irrelevant task template (average pattern-similarity = 0.08, CI = [0.06, 0.10], bootstrap *p* < .001 Fig. 2B).

**Figure 2.**
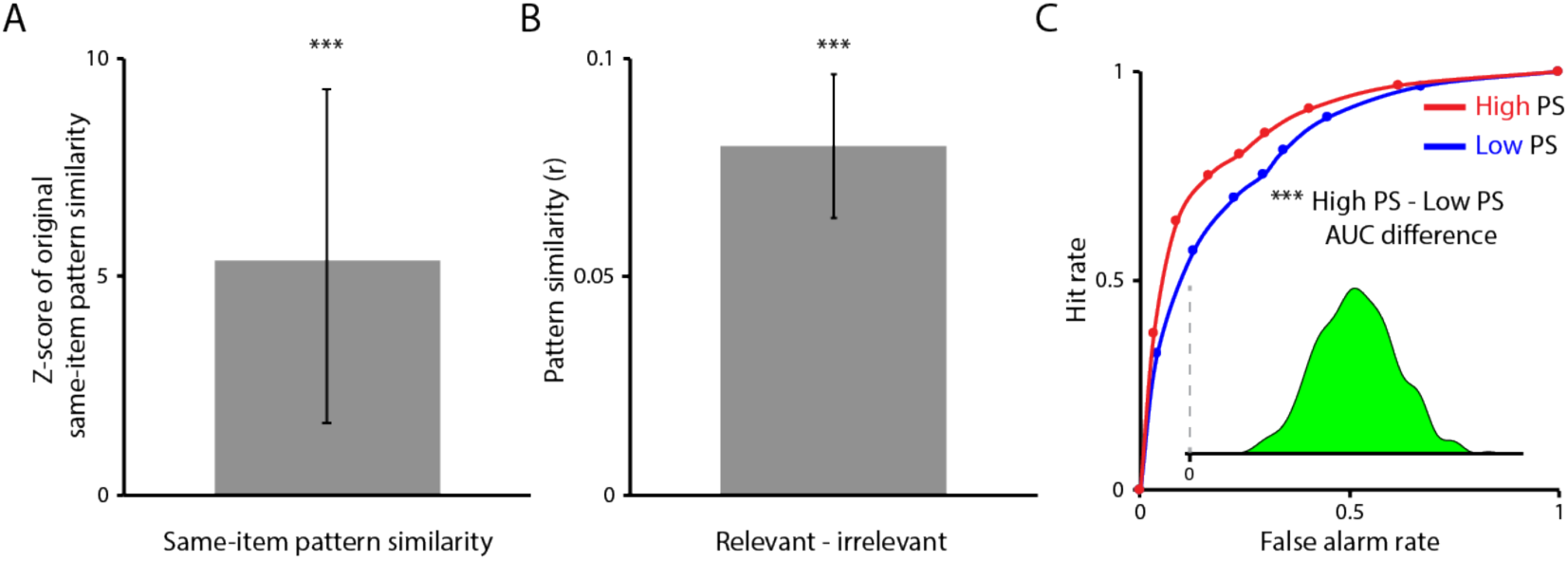
Manipulation checks. (A) Same-item pattern similarity in LOC was higher than pattern similarity scores based on permuted pairings. (B) During the first phase, pattern similarity was higher for the relevant (current) task than the irrelevant task. (C) Greater evidence for the current task’s template during encoding was associated with better task memory on the final test, as reflected in a greater AUC for high vs. low pattern similarity. *** *p* < .001.

### Task encoding results

Several previous studies (Gordon, Rissman, Kiani, & Wagner, 2014; Kim et al., 2014; Koen & Rugg, 2016; Kuhl, Rissman, & Wagner, 2012) have reported positive relationships between multivariate measures of encoding strength and subsequent memory. Consistent with this, greater neural evidence for the corresponding task template during encoding was associated with better task memory on the final test (AUC difference = 0.05, CI = [0.03, 0.07], bootstrap *p* < .001; Fig. 2C).

### Relationship between item reactivation and subsequent source memory

We hypothesized that reactivation of the item representation from the first phase during the second phase would lead to retroactive interference with the source memory for the initial task. Consistent with our hypothesis, higher pattern similarity between the first and second phases was associated with worse source memory (AUC difference = -0.04, CI = [-0.07, -0.01], bootstrap *p* = .002; Fig. 3C).

**Figure 3.**
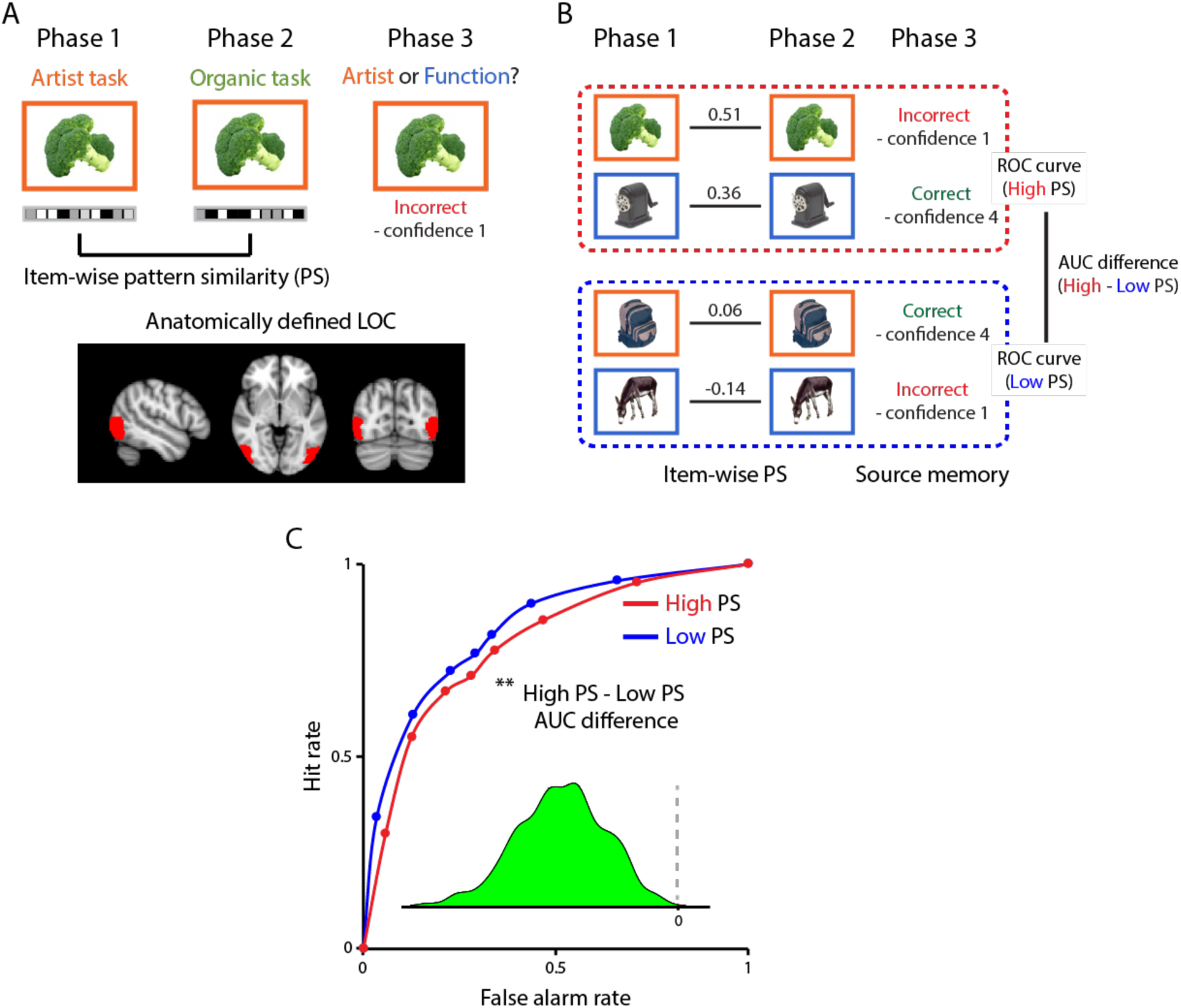
Relating pattern similarity to source memory across items. (A) Pattern similarity was measured for each item between the first and second phases. (B) After performing a median split of the pattern similarity scores, an ROC curve was constructed for high and low pattern similarity items from the source memory test and the AUC was calculated. A difference in these AUCs indicates a relationship between pattern similarity and source memory, and the sign indicates the direction of the relationship. (C) There was a reliable negative relationship between pattern similarity and source memory across items. ** *p* < .01

This observed negative relationship might be driven by item-wise variance in task reactivation rather than item reactivation. We controlled for this confound using a two-step permutation analysis (see Methods): First, we regressed out generic task information from patterns in the first and second phases (Fig. 4A), and re-calculated pattern similarity and its relationship to source memory. Second, we scrambled the original pairings of items across the first and second phases and re-calculated the relationship for each permutation (Fig. 4C). If item reactivation modulates subsequent source memory, as hypothesized, then the relationship based on residual intact pairings (excluding task information) should be more negative than the null distribution of relationships from permuted parings (excluding both task and item information).

**Figure 4.**
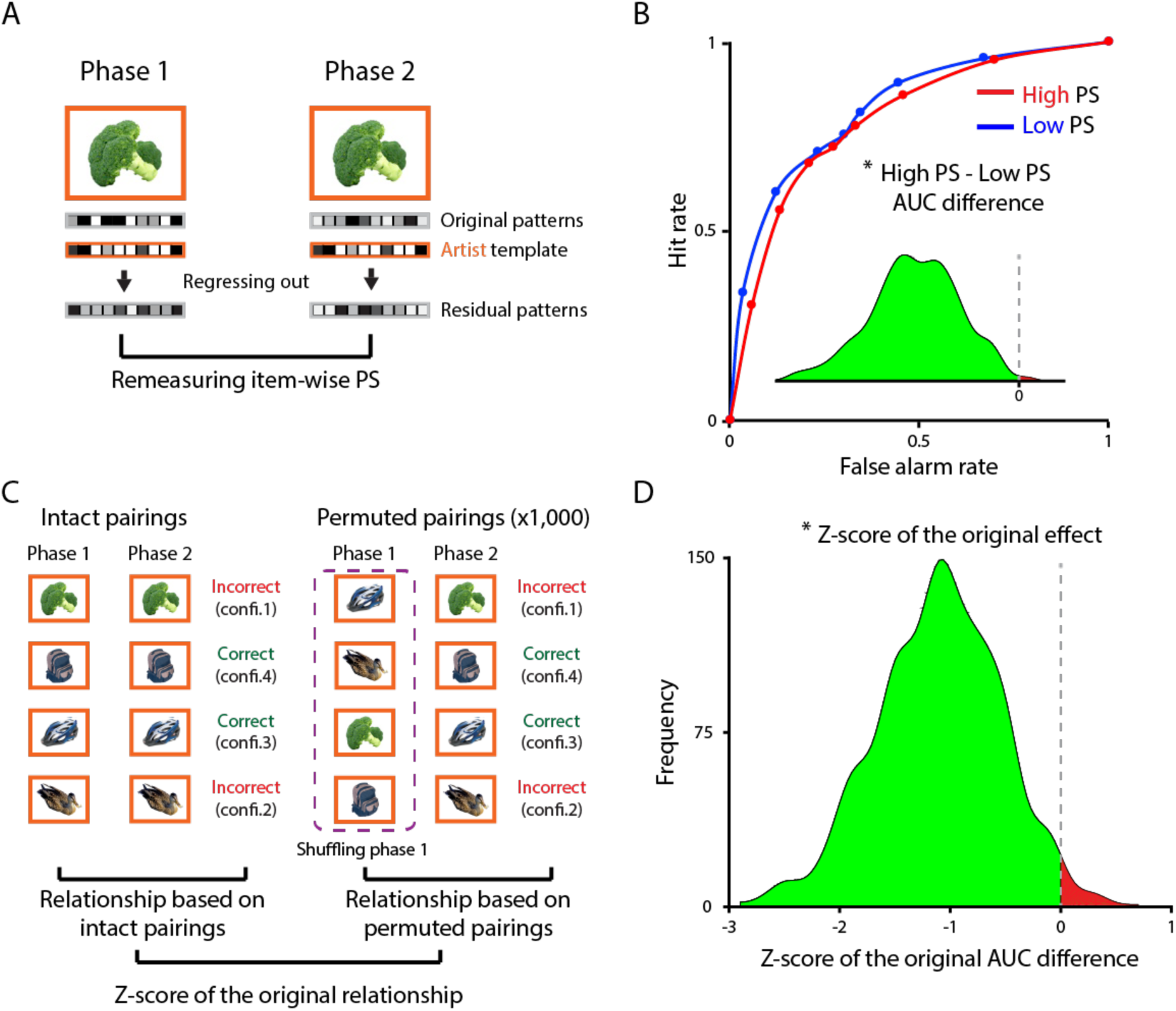
Testing specificity of item reinstatement effects. (A) Generic task information was regressed out of the item patterns from the first and second phase using the task templates from the localizer. (B) Using the residual patterns, the relationship between item-wise pattern similarity and source memory was re-calculated and remained reliably negative. (C) We further narrowed in on item-specific variance by submitting the residual item patterns to a permutation test. (D) The *z*-score of the original negative relationship was more negative that the null relationships calculated after permuting the item pairings between the first and second phases. * *p* < .05.

After the first step of regressing out task information, the relationship for intact pairings remained negative (AUC difference = -0.03, CI = [-0.06, -0.01], bootstrap *p* = .020; Fig. 4B). In addition, this relationship was significantly more negative than the null distribution of permuted relationships after the second step (average *z*-score = -1.10, CI = [-2.28, -0.04], bootstrap *p* = .042; Fig. 4D).

### Relationship between task reactivation and subsequent source memory

In addition to examining the effect of item reinstatement on initial source memory, we also considered the effect of task reinstatement. We measured the amount of information about the initial task when each item was repeated in the second phase by calculating pattern similarity with the task templates from the localizer (Fig. 5A and 5B). In contrast to the negative effect of item reinstatement, there was a numerically positive effect of task reinstatement, although it failed to reach significance (AUC difference = 0.02, CI = [-0.01, 0.05], bootstrap *p* = .21; Fig. 5C).

**Figure 5.**
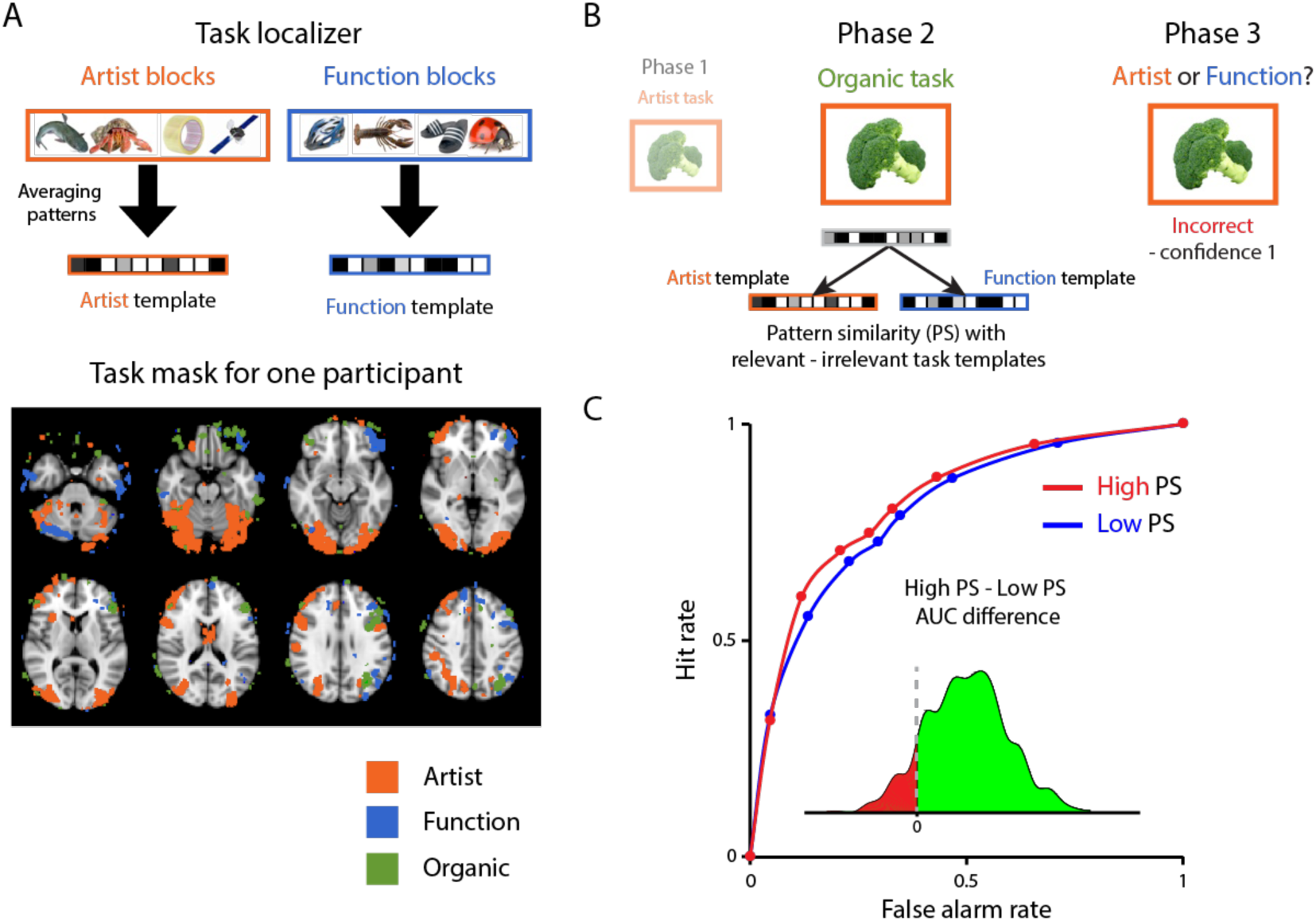
Relating task reactivation to memory. (A) We generated template activity patterns for each task from the localizer by selecting task-selective voxels with a GLM and then averaging the activity values in each voxel across all trials of the task. (B) We measured reactivation of the initial task (artist or function) in the second phase as the pattern similarity of each item with its corresponding task template minus its non-corresponding task template (both of which were different from the current organic task). (C) The relationship between this index of task reactivation and subsequent source memory was numerically positive, but not significant.

### Ruling out univariate confounds

We have assumed that item-wise pattern similarity reflects neural overlap of an item representation across the first and second phases, and that this neural overlap affects subsequent retrieval of the initial task. In principle, however, univariate activation in the second phase might have affected both pattern similarity and subsequent source memory. For example, imagine that a participant was in an inattentive state for some of trials in the second phase. Lower activation for those trials (vs. more attentive trials) could reduce item-wise pattern similarity across the first and second phases (Coutanche, 2013; Davis and Poldrack, 2013; Davis et al., 2014; Aly and Turk-Browne, 2016). Furthermore, subsequent source memory for the initial task on those inattentive trials might be better, because there was less learning of second-phase information, leading to less retroactive interference. In short, univariate activation in the second phase might be a factor determining the negative relationship between item-wise pattern similarity and subsequent source memory.

Importantly, the item-specificity of the effects argues against this explanation: If the negative relationship between item-wise pattern similarity and subsequent memory solely depended on univariate activation level in the second phase, then the same effect should have persisted even after permuting the original item pairings (Figure 4C and 4D). Despite the implausibility of this explanation, we ran an analysis to test it; specifically, we computed the relationship between univariate LOC activation in the first and second phases and subsequent source memory, using the same analysis procedures as for pattern similarity. First-phase and second-phase univariate activation were not reliably related to subsequent memory (*p*s > .10). The difference in univariate activation between the two phases (phase 1 minus phase 2) failed to predict memory as well (*p* = .23). These findings are consistent with our interpretation that the negative relationship between item-wise pattern similarity and subsequent source memory reflects neural overlap of items across the two phases rather than overall activity.

## Discussion

We investigated how memory for the context in which an object was encountered is influenced by encountering it again in a novel context. Although the basic behavioral finding — impaired subsequent source memory for the initial context — has been demonstrated previously (McGovern, 1964; Postman & Underwood, 1973; Richter et al., 2016), we tested a novel explanation for this important phenomenon. Specifically, we hypothesized that the extent to which the item is represented the same way across contexts determines how much interference the old context suffers. Consistent with this hypothesis, we found a negative relationship between neural overlap across item repetitions and subsequent source memory for the initial context.

We used pattern similarity to index how much the initial item representation was reactivated upon repetition. However, taken at face value, this measure does not necessarily reflect item information alone — it can also be influenced by variance in task reactivation. We addressed this issue by showing: (a) that the negative relationship persists after regressing out task information, (b) that this relationship is eliminated after permuting item pairings within task, and (c) that task reactivation *per se* leads to an effect that (numerically) goes in the opposite direction. Taken together, these results strongly support our main conclusion that reactivation of an item-specific representation leads to retroactive interference.

Using a paradigm similar to ours, Koen and Rugg (2016) also recently investigated the consequence of item repetition for task memory. As in our study, participants performed one task on an item during an initial phase and then performed a different task on that item during a later phase; participants were then required to recall both tasks for each item. Interestingly, Koen and Rugg (2016) found a *positive* relationship between reactivation during item repetition and subsequent source memory for the initial task — the opposite of what we found.

However, we think that these results are not necessarily incompatible with ours, given several differences between the studies. Most importantly, we speculate that the contents of reactivation might have been different. Our use of object images (their materials were words) allowed us to concentrate analyses on LOC, a region known to represent item-level features of objects (Cichy et al., 2011; Eger et al., 2007). In contrast, Koen and Rugg (2016) measured pattern similarity from a task-selective mask, which by design likely contained more task information than item information. Thus, it is possible that item-related pattern similarity in their study reflected initial task information specific to each item. For example, after making a pleasantness judgment (an initial task in their study) about a word like “broccoli”, later reactivation might contain task information specific to that item, such as that it has an unpleasant taste. This retrieved information related to the initial task is unlikely to be confused with the second task in the final source memory test because the tasks are highly distinct from each other. In fact, reactivation of this task information might strengthen its association with the item and ultimately improve source memory.

In our case, because we focused on a region representing item-specific information, pattern similarity was more likely to reflect how reliably the component *shared* across the repeated experiences — i.e., the item — was represented. Reliable reinstatement of the initial item representation can lead that representation to also become associated with the second task, which later interferes with retrieval of the initial task (Gershman, Schapiro, Hupbach, & Norman, 2013; Hupbach, Gomez, Hardt, & Nadel, 2007; St. Jacques, Olm, & Schacter, 2013; Sederberg, Gershman, Polyn, & Norman, 2011). Put simply, we speculate that the relationship between item pattern similarity and source memory might depend on the contents of the measure: item pattern similarity related to idiosyncratic task information can enhance source memory, whereas item pattern similarity related to specific item features can lead to retroactive interference.

Generally speaking, these results highlight the complex nature of retroactive interference effects. As prior work has shown, there is no simple answer to the question of how new learning modulates the accessibility of existing knowledge: There is some evidence that reactivation makes old memories susceptible to retroactive interference (Forcato et al., 2007; Gershman et al., 2013; Hupbach et al.,; Sederberg et al., 2011), but other studies observed the opposite effect that reactivation alleviates retroactive interference (Kuhl et al., 2010; Koen and Rugg, 2016). Still others posit that the *degree* to which information is activated determines whether it is strengthened or weakened (e.g., Detre et al., 2013). Our findings contribute to this debate by suggesting that — when an item appears in multiple contexts — the specific *content* of the reactivated memory is a key determinant of retroactive interference: If contextual features uniquely related to the item in the initial experience are strongly reactivated during new learning, this can strengthen memory for these features (e.g., Koen and Rugg, 2016; Kuhl et al., 2010; a trend in our study). However, crucially, when shared *item* features are reactivated, our results show that this can impair memory for the initial context.

## Supplementary Materials

### Simulations: Testing validity of analyses

As noted in the main paper, our central prediction concerns the relationship between reactivation of an item’s representation and subsequent source memory. To test this, we needed a way of measuring reactivation of item features and relating this to memory performance. Our basic approach to this problem was to compute pattern similarity in LOC across the two presentations of an item (in phase 1 and phase 2), and then correlate this measure with source memory for the initial task (e.g., artist task in phase 1). However, this analysis is complicated by the fact that LOC pattern similarity is potentially influenced by two factors — reactivation of item features and reactivation of generic artist or function task features — which could influence memory in different (potentially opposing) ways. For this reason, it is essential to use an analysis procedure that properly separates the effects of item vs. task reactivation on subsequent memory.

To assist us in making this determination, we ran simulations contrasting two situations: (A) where subsequent memory was affected by item-feature reactivation but not generic task reactivation (we will call this the *item reactivation* simulation, illustrated in Figure S1A); and (B) where subsequent memory was affected by generic task reactivation but not item reactivation (we will call this the *task reactivation* simulation, illustrated in Figure S1B). We explored how well different candidate analysis procedures (used in previous studies by Koen & Rugg, 2016, and by Kim et al., 2014) could separate out these influences. Our goal was to find an analysis procedure that reports a positive result *only* in the first situation (i.e., item-feature reactivation is driving subsequent memory but not task reactivation).

**Figure S1.**
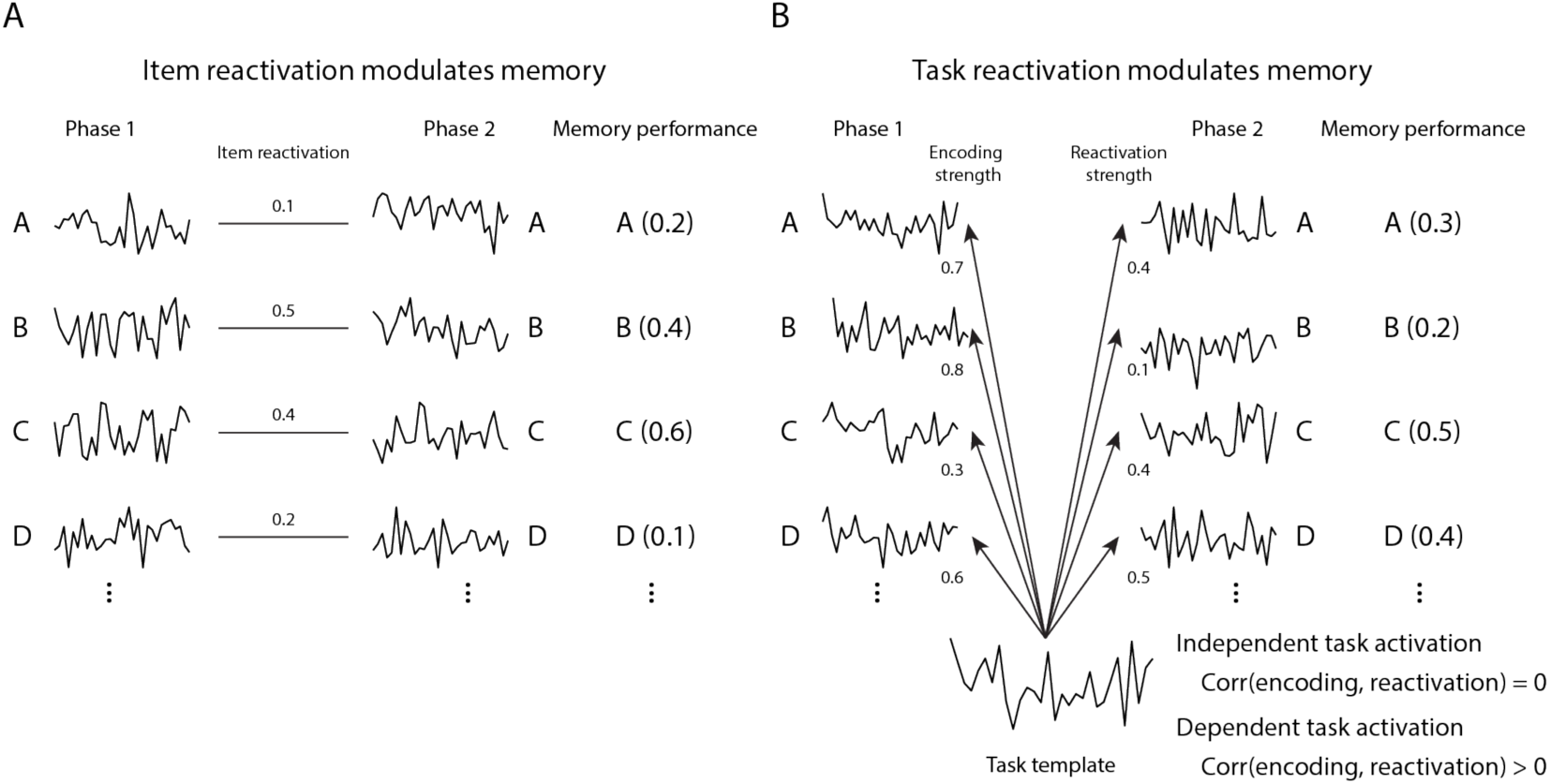
Generating simulated data. (A) In the *item reactivation* simulations, phase 1 patterns were drawn randomly for each item, and an item reactivation score was drawn randomly for each item. Phase 2 patterns were generated by adding noise to corresponding phase 1 patterns, such that the correlation between phase 1 and phase 2 patterns (for a given item) was equal to that item’s reactivation score. Memory performance scores were also generated based on item reactivation scores, such that the two variables were negatively correlated across items. (2) In the *task reactivation* simulations, we randomly generated a task template, and *encoding strength* and *reactivation strength* scores were drawn randomly for each item. Phase 1 and phase 2 patterns were generated by adding noise to this task template, such that the correlation between the task template and the phase 1 pattern for an item was equal to that item’s encoding strength, and the correlation between the task template and the phase 2 pattern for an item was equal to that item’s retrieval strength. For the independent task activation simulation variant, encoding and reactivation strength were independent, and for the dependent task activation variant, the two variables are positively correlated (i.e., stronger encoding 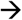 greater reactivation). Memory strength was generated based on reactivation strength, such that the two variables were negatively correlated across items.

#### Generating simulated data

For each simulation, we generated two sets of 100 patterns, each containing 1,000 features assigned a value sampled from a uniform distribution between 0 and 1. The first and second sets of patterns correspond to the encoding phase (phase 1) and repetition phase (phase 2), respectively, and each pattern corresponds to the (simulated) activation pattern evoked when an item is presented.

In the first type of simulation, where *item reactivation* determines activation patterns and memory performance, we randomly generated 100 patterns in phase 1. Each item was also assigned an item reactivation score (mean = 0.3, SD = 0.1; Figure S1A). Phase 2 patterns were generated by adding zero-mean Gaussian noise to phase 1 patterns, such that the correlation between the phase 1 and 2 patterns for a given item was equal to that item’s reactivation score. Finally, we also generated a memory performance score for each item based on the item reactivation score, so that the two variables were *negatively* correlated (mean correlation coefficient = -0.3, SD = 0.1).

For the second type of simulation, where *task reactivation* determines memory, we randomly generated a task template, and also an *encoding strength* and *reactivation strength* score for each item. Next, we generated phase 1 and 2 patterns by adding zero-mean Gaussian noise to the task template, such that the correlation between the task template and each item’s phase 1 pattern was equal to that item’s encoding strength score, and the correlation between the task template and each item’s phase 2 pattern was equal to that item’s retrieval strength score.

We ran two subtypes of the *task reactivation* simulation: In the first variant (*independent task activation*), encoding and reactivation strength were determined independently; in the second variant (*dependent task activation*), task encoding and reactivation strength were positively correlated (such that stronger task encoding leads to greater task reactivation; mean correlation coefficient = 0.3, SD = 0.1). First, we randomly generated a task template, and patterns in phase 1 and phase 2 were generated based on task encoding strength (phase 1) and reactivation strength (phase 2) respectively (Figure S1B). For example, for a given item, if task encoding strength is 0.7, a pattern in phase 1 was generated so that correlation coefficient with a task template to be 0.7. On average, encoding strength was greater than reactivation strength (mean encoding strength = 0.6, mean reactivation strength = 0.3, SDs = 0.1). For both variants of the task reactivation simulation, we generated a memory performance score for each item that was determined by task reactivation strength, such that the two variables were *negatively* correlated (mean correlation coefficient = -0.3, SD = 0.1).

For each simulation type, we ran 1,000 simulations (each simulation corresponds to a single participant), and we tried multiple analysis methods on each dataset. As stated above, our goal was to find an analysis procedure that *only* returns a significant result if there is a relationship between item reactivation and subsequent memory performance. Put concretely, we were seeking an analysis that returns a significant result for the *item reactivation* simulation type but not for the *task reactivation* simulation type.

#### Same minus different item pattern similarity

The first analysis type was based on the procedures used in Koen & Rugg (2016), who computed the difference between *same-item pattern similarity* (i.e., the similarity between an item’s phase 1 pattern and the same item’s phase 2 pattern) and *different-item pattern similarity* (i.e., the similarity between phase 1 and 2 patterns corresponding to different items) and then related this to memory performance. First, we calculated *same-item pattern similarity* scores by calculating the Pearson correlation coefficient of items across phase 1 and 2 (e.g., A in phase 1 to A in phase 2). Second, we measured average pattern similarity score of items in phase 1 with different items in phase 2 (e.g., A in phase 1 to B, C, D in phase 2), and did this calculation in the opposite direction (e.g., A in phase 2 to B, C, D in phase 1), and averaged these two to generate *different-item pattern similarity* scores. We then subtracted different-item pattern similarity from same-item pattern similarity score for each item. Finally, we measured the correlation between this same minus different item pattern similarity score and memory performance across items.

We performed this analysis for each of simulation types (i.e., item reactivation; task reactivation with independent task activation; task reactivation with dependent task activation). We observed that the relationship was negative for every simulation type (*ps* < .001) — that is, this analysis method yielded significant results both when memory was driven by item reactivation *and* when memory was driven by task reactivation (but not item reactivation), making it unsuitable for our present purposes.

Why does this analysis find significant results when memory is driven by task reactivation? One might think that comparing same-item pattern similarity to different-item pattern similarity (when all items were encoded with the same task) would control for task reactivation. However, crucially, the method used here for computing different-item pattern similarity does not perfectly cancel out the effects of task reactivation. Because patterns in the task reactivation simulations are generated from a common task template, same-item pattern similarity can be approximately estimated by multiplying encoding (x) and reactivation (y) strength (same ≈ *xy*). Given that memory performance (by design) was correlated with task reactivation (*y*) in this simulation, it stands to reason that same-item pattern similarity (≈ *xy*) and memory performance will be correlated (they share an influence of y). While different-item pattern similarity (for a given item A) is influenced by A’s reactivation strength, it is also influenced by other terms that have nothing to do with A’s reactivation strength (e.g., the similarity of the phase 1 A pattern and the phase 2 B pattern). The key issue is that these other influences dilute the influence of A’s reactivation strength and prevent the different-item pattern similarity term from perfectly canceling out the same-item pattern similarity term.

#### Permutation analysis

The next analysis method we tested was permutation analysis, which was used in our previous study (Kim et al., 2014). First, we removed item-specific information from pattern similarity scores by permuting the original item pairings. Specifically, we shuffled the phase 1 items 1,000 times while keeping the phase 2 ordering the same (e.g., A-A, B-B, C-C, D-D 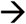 B-A, C-B, D-C, A-D). Then, for each permutation, we re-measured the relationship between item-wise pattern similarity and memory (Figure S2A). This procedure gives us a null distribution of 1,000 correlation coefficients. As the final step, we computed the *z*-score of the original correlation coefficient (based on the intact pairings) relative to the null distribution (Figure S2B).

**Figure S2.**
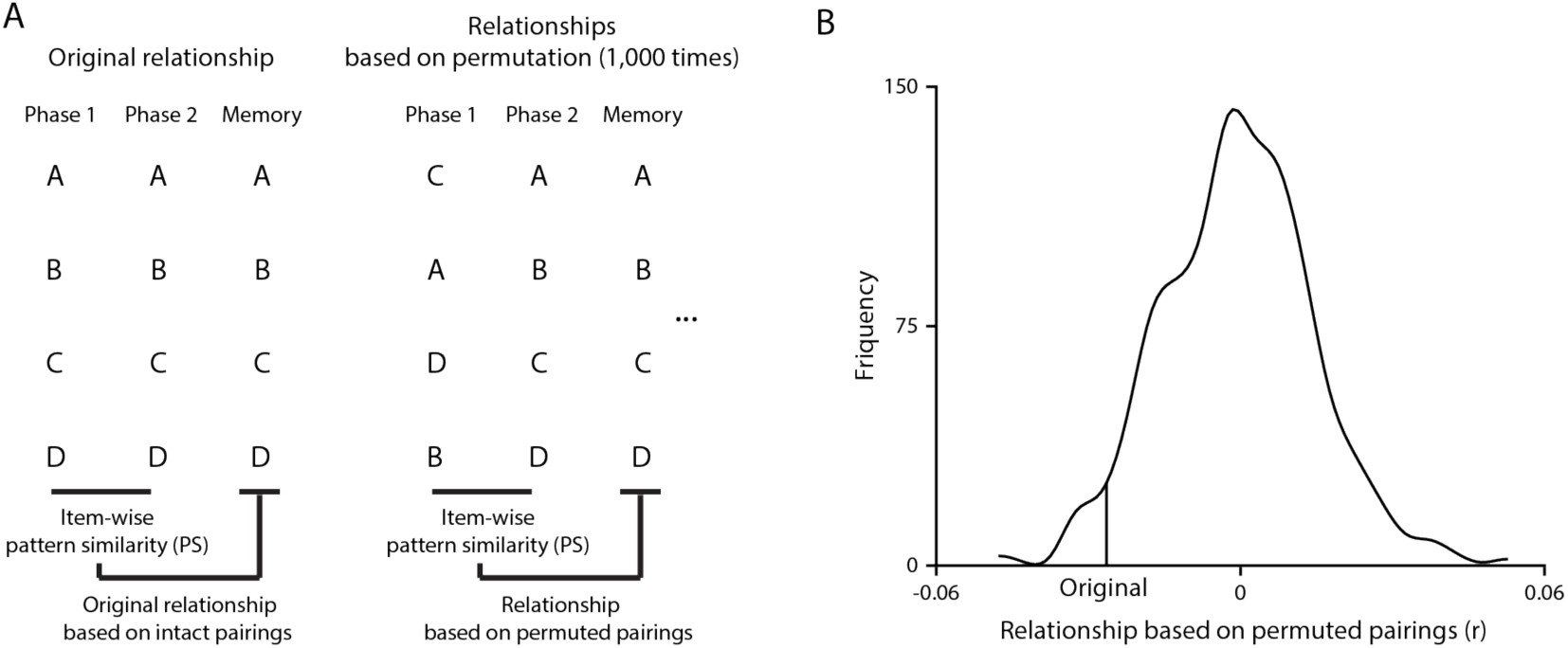
Permutation analysis. (A) First, based on intact item parings (across phase 1 and 2), we measured the original relationship between item-wise pattern similarity and memory. Second, we permuted the item pairings (by shuffling phase 1 items) 1,000 times and measured the relationship between item-wise pattern similarity and memory for each of permutation. (B) We calculated the *z*-score of the original relationship (from intact pairings) based on the null distribution of 1,000 relationships acquired from the permutations.

For the *item reactivation* simulation type, the original negative relationship was significantly more negative compared to the relationships acquired from permutation (mean *z*-score = -2.91, *p* < .001). In contrast, for the *task reactivation* simulation type with *independent task activation* (i.e., where, for a given item, the presence of the task pattern at encoding was independent from the presence of that pattern at retrieval), the original relationship was not different from those of permutation (mean *z*-score = -0.02, *p* = .53). Thus, assuming that task encoding and reactivation are independent, the permutation analysis approach achieves our goal of returning a significant result when subsequent memory is linked to item reactivation, but not when subsequent memory is linked to task reactivation.

However, problems arise when we consider the third type of simulation: task reactivation with *dependent task activation* (i.e., where, the presence of the task pattern at encoding was correlated with the presence of that pattern at retrieval). For this simulation, the permutation analysis procedure returns a significant result — that is, the original negative relationship was significantly more negative compared to the relationships acquired from permutation (mean *z*-score = -0.13, *p* < .001). This occurs because the correlation between encoding and reactivation strength provides additional pair-wise variance (correlated with task reactivation) that is removed by permutation, resulting in a difference between the original and permuted results.

#### Regressing out generic task information

As demonstrated above, the permutation analysis fails to meet our goal (of returning significant results *only if* subsequent memory is linked to item reactivation) in the case where task encoding and reactivation strength are correlated. To address this remaining issue, we explored a new procedure where we regressed out generic task information from patterns in phase 1 and 2 and *then* ran the permutation. Specifically, we ran a regression where we predicted each pattern (in phase 1 and phase 2) using the task templates from the localizer, and we took the residuals from this analysis; based on the residual patterns, we performed the permutation analysis again.

We expected that this new analysis would (properly) return a null result for the task reactivation simulation (dependent task activation variant) because the regression should remove any item-wise pattern similarity caused by task reactivation. The results of the analysis supported our predictions: the original correlation was not significantly different from the permuted distribution (mean *z*-score = 0.03, *p* = .37). The same results held for the independent task activation variant too (mean *z*-score = -0.01, *p* = .69). This new analysis successfully distinguished the original correlation from the permuted distribution (mean *z*-score = -2.95, *p* < .001) only when item reactivation determines memory (i.e., *item reactivation* simulation type).

In sum, this final analysis approach (regressing out task patterns, followed by permutation) meets our full set of desiderata, returning a significant result for the item reactivation simulation type but neither of the task reactivation simulation types. Based on these results, this is the approach that we decided to use in the paper.

Kim G, Lewis-Peacock JA, Norman KA, Turk-Browne NB. 2014. Pruning of memories by context-based prediction error. Proc Natl Acad Sci. 111:8997–9002.

Koen JD, Rugg MD. 2016. Memory reactivation predicts resistance to retroactive interference: Evidence from multivariate classification and pattern similarity analyses. J Neurosci. 36:4389–4399.

